# Estimating Movement Smoothness from Inertial Measurement Units

**DOI:** 10.1101/2020.04.30.069930

**Authors:** Alejandro Melendez-Calderon, Camila Shirota, Sivakumar Balasubramanian

## Abstract

There is an increasing trend in using inertial measurement units (IMUs) to estimate movement quality and quantity, and infer the nature of motor behavior. The current literature contains several attempts to estimate movement smoothness using data from IMUs, most of which assume that the translational and rotational kinematics measured by IMUs can be directly used with existing smoothness measures - spectral arc length (SPARC) and log dimensionless jerk (LDLJ-V). However, there has been no investigation of the validity of these approaches. In this paper, we systematically evaluate the appropriateness of the using these measures on the kinematics measured by an IMU. We show that: (a) current measures (SPARC and LDLJ-V) are inappropriate for translational movements; and (b) SPARC and LDLJ-V can be used rotational kinematics measured by an IMU. For discrete translational movements, we propose a modified version of the LDLJ-V measure, which can be applied to acceleration data (LDLJ-A), while roughly maintaining the properties of the original measure. However, accuracy of LDLJ-A depends on the IMU orientation estimation error. We evaluate the performance of these measures using simulated and experimental data. We then provide recommendations for how to appropriately apply these measures in practice, and the various factors to be aware of when performing smoothness analysis using IMU data.

## 1 Introduction

Inertial Measurement Units (IMUs) are becoming ubiquitous in everyday objects we carry, e.g. smartphones, smart watches, smart clothing, etc. This, along with the fact that IMUs are now relatively inexpensive, has sparked their use for movement analysis in different disciplines, such as movement neuroscience (Shull et al., 2014; Picerno, 2017; O’Reilly et al., 2018), movement biomechanics and sports science (Salmond et al., 2017; Li et al., 2016; Johnston et al., 2019), and neurorehabilitation (Brognara et al., 2019; Dobkin, 2013; Hubble et al., 2015; Parker et al., 2020; Vienne et al., 2017; Wang et al., 2017).

Quantitative measures of *movement smoothness* are of great interest as they allow us to evaluate the evolution of motor skill learning or recovery (Trehan et al., 2015; Moghaddas et al., 2019; Toosizadeh et al., 2015). While there have been attempts to formally define *movement smoothness* (Hogan and Sternad, 2007, 2009; Balasubramanian et al., 2012, 2015), there is still no consensus on the most appropriate measure to use in different tasks or with different measurement technologies. This has led to the adoption of a diverse set of smoothness measures, which can often lead to contradicting results on the same data, leading to inconsistent and potentially incorrect interpretations (Fig. 1).

**Figure 1:**
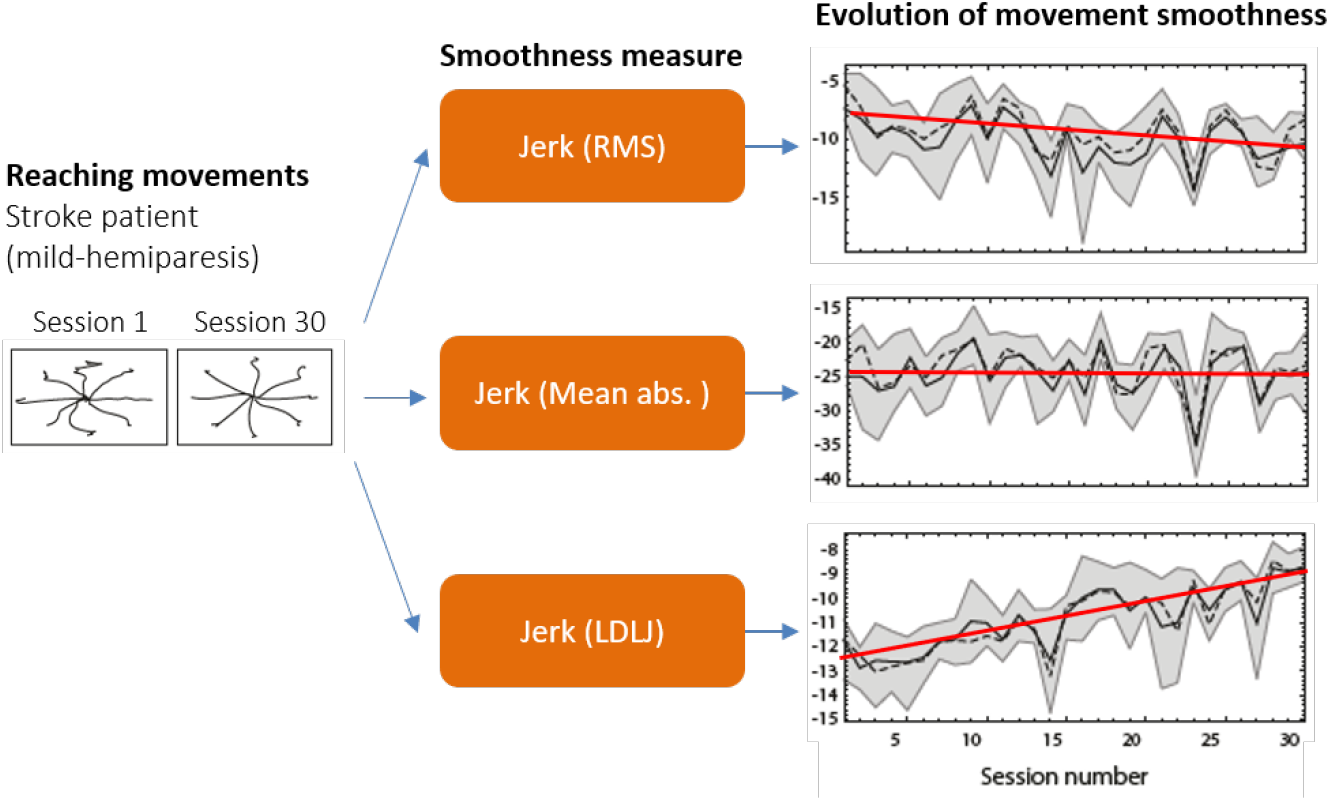
Different measures of smoothness can lead to vastly different conclusions. In this example, three jerk-based smoothness measures (middle) were applied to arm movements performed by a post-stroke subject in multiple directions (left), over 30 training sessions (right). The trends suggest three conflicting outcomes: Jerk (RMS) shows reduction in movement smoothness, indicating that the training was detrimental; Jerk (Mean abs.) indicates that the training had no effect; and Jerk (LDLJ) indicates that the training was beneficial. Data from (Balasubramanian et al., 2012) with permission.

To ensure appropriate quantification and interpretation, valid, consistent, and reliable measures of *movement smoothness* should be used. Examples of such measures are the SPARC (spectral arc length) and LDLJ (log dimensionless jerk) applied to the movement’s velocity (Balasubramanian et al., 2012, 2015). Since their proposal, these measures have been used in multiple studies using data from different motion sensing technologies (e.g., (Gulde and Hermsdörfer, 2018; Colombo et al., 2017; Rihar et al., 2014; Abdi et al., 2016)). In many cases, especially when movement velocity is not directly available, the SPARC or LDLJ have been loosely adapted to other kinematic variables, e.g., acceleration from IMUs. Possible adaptations of these measures for different signals were tentatively proposed in Balasubramanian et al. (2015); however, they were not supported by any mathematical arguments or empirical data. Thus, the properties of these smoothness measure adaptations for different signals are unknown. Further, it is likely that the smoothness values derived from different variables (e.g., velocity or acceleration) are not directly comparable and may even lead to different interpretations (e.g., Fig. 1).

In this paper, we present a systematic investigation of the use of SPARC and LDLJ to evaluate movement smoothness using IMU data – linear acceleration from the accelerometer and angular velocity from the gyroscope. We show that the SPARC cannot be used with IMU (accelerometer) data to estimate smoothness of translational movements, and propose a modification to the LDLJ measure for this purpose, while maintaining the measure’s properties. For rotational movements, we show that the SPARC can be used without any changes with gyroscope data, while LDLJ is prone to errors. Using simulated and experimental data, we show conditions under which movement smoothness analysis can be carried out on both translational and rotational movements using an IMU. We strongly believe that the methods proposed in this paper are critical to (a) standardize analysis methods used in similar contexts (e.g. movement type, measurement technology, etc.); (b) avoid biased or inappropriate selection of smoothness measures; and (c) facilitate interpretation and direct comparison of results between studies.

## 2 Movement smoothness from IMUs: preliminaries

In this section, we present two main problems associated with the use of IMUs to analyse movement smoothness:

1. the construct of movement smoothness: why velocity cannot simply be replaced by acceleration in the existing smoothness “formulae”;
2. the kinematics measured by IMUs: transforming IMU measurements from local to world reference frames are prone to errors.

### 2.1 The construct of *movement smoothness*

Intuitively, smoothness is an intrinsic property of a movement, and should not depend on the variables used to represent that movement. That is, the smoothness value obtained from two different representations of a movement *M* (e.g., its velocity **v**(*t*) or acceleration **a**(*t*)) should quantify the same underlying construct of *movement smoothness*. Thus, given that existing smoothness measures are defined from velocity (see equations below), if a movement’s velocity were not directly accessible–such as in the case of an IMU–a makeshift solution could be to simply replace the velocity terms with acceleration to compute movement smoothness.

Although this approach is simple and appealing–and used in several research papers (e.g., (Dixon et al., 2018; Beck et al., 2018)), it is semantically incorrect as a measure of the smoothness of the movement represented by the acceleration signal. Almost all smoothness measures have been defined using velocity as the starting point; this is probably because velocity provides a good representation of movement intermittency, and thus acts as a reasonable reference for developing smoothness measures. The SPARC was developed using submovements as a putative model of movement generation, resulting in the definition in Eq. 1 that quantifies the dispersion of submovements in time. On the other hand, the LDJL uses the minimum jerk model of movement planning, which results in the definition in Eq. 2.

#### SPARC

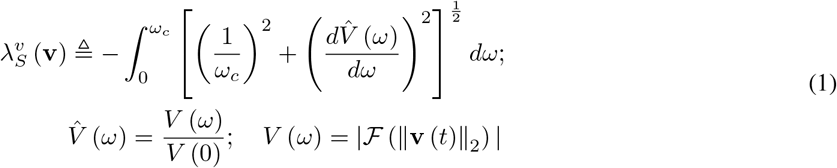

#### LDLJ (based on velocity; LDLJ-V)

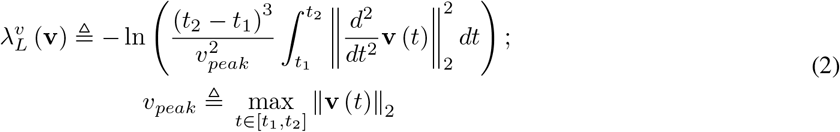

where **v**(*t*) represents the velocity (linear or angular) of a movement in the time domain, *V* (*ω*) is the magnitude of the Fourier transform 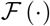 of the movement speed, *ω_c_* is an adaptive cut-off frequency (see (Balasubramanian et al., 2015) for details), and *t*_1_ and *t*_2_ the start and stop times of the movement. It should be noted that we will refer to the LDLJ measure in Eq. 2 as LDLJ-V in the rest of the manuscript, to distinguish it from the LDLJ-A measure introduced later (Eq. 15).

From a movement analysis perspective, SPARC and LDLJ-V intrinsically perform a shape analysis operation, assigning values to the shape of a movement’s velocity profile. Properties that are characteristic of these two measures, such as monotonic responses to the number of submovements and inter-submovement intervals, may not remain when replacing velocity of a movement with its acceleration in the definitions above. Indeed, we show in Appendix B that directly replacing velocity by acceleration in Eq. 1 and Eq. 2 results in undesirable properties that fog the interpretation of the smoothness values.

Thus, it is crucial to be aware of the type of signal (position, velocity, acceleration, etc.) available for smoothness analysis and ensure that it is consistent with the definition of movement smoothness adopted in each application. To ensure a consistent approach, SPARC and LDJL-V should be computed only from movement velocity. When velocity is not directly available, as in the case of IMU data from translational movements, the following sections provide recommendations.

### 2.2 The kinematics measured by a single IMU

To better understand the problems associated with the use of a single IMU data for the analysis of *movement smoothness*, consider an IMU undergoing translational and rotational movements (Fig. 2). An IMU typically includes a 3-axis accelerometer, which measures its linear acceleration 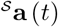, and a 3-axis gyroscope, which measures its angular velocity 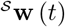. Both measurements are with respect to 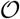 (an inertial reference frame) expressed in the IMU’s local (sensor) reference frame 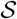:

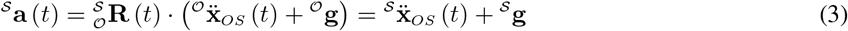

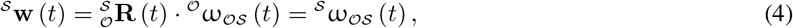

where 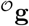 is the gravity vector, and 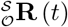 is the rotation matrix representing 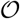 in 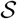. For simplicity, we will drop the notation indicating the dependence on time *t* for the different variables (e.g. 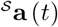 will be written as 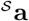). In the analysis presented here, we do not consider the contributions of measurement noise, as our primary goal is to understand the effect of the physics of IMUs on smoothness analysis. Some work on the effect of noise on smoothness analysis has been presented in our earlier work (Balasubramanian et al., 2012, 2015).

**Figure 2:**
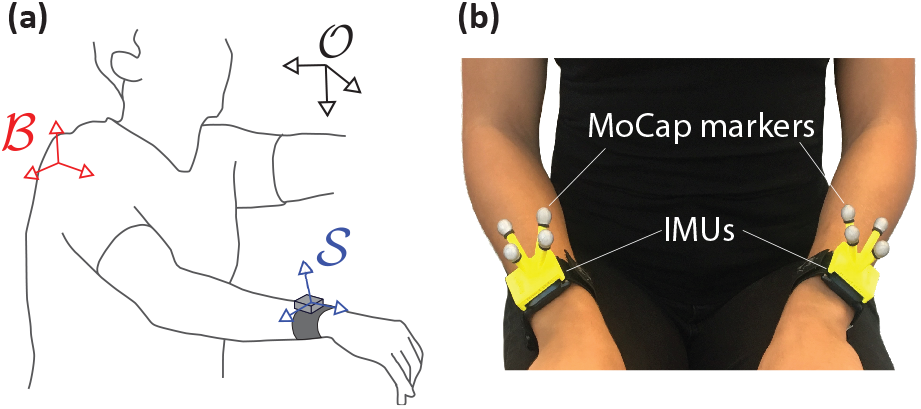
(a) Different coordinate frames of interest in movement analysis: inertial or earth-fixed 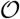, body 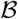, and sensor 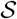 reference frames. (b) Different technologies used to capture the kinematics of 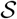: motion capture using reflective markers (MoCap) or Inertial Motion Units (IMUs). This setup was used to collect the data presented in section 4.2.

In human movement analysis, we are typically interested in translational and rotational motions of a single or a chain of body segments relative to a body coordinate frame 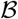, while measurements are often made with respect to an earth-fixed coordinate system 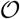 (assumed to be an inertial reference frame). Consider the scenario depicted in Fig. 2, where we are interested in the translational and rotational movements of the wrist 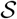, with respect to the shoulder reference frame 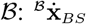 and 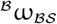, respectively. It should be noted that 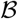 is physically attached to 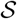 through a set of rigid bodies interconnected through joints, and thus any translational or rotational movement of 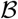 will be transmitted to 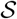, as detailed below.

#### Translational motion

In the example in Fig. 2, the accelerometer signal 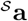 is composed of (derivation in Appendix A):

1. linear acceleration due to gravity 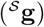;
2. linear acceleration of 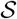 w.r.t. 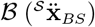;
3. linear acceleration of 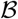 w.r.t 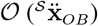;
4. Coriolis acceleration of 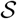 due to rotation of frame 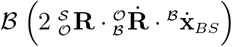; and
5. linear acceleration of 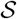 due to angular acceleration of frame 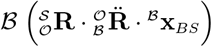.

Thus, in general, 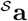 cannot be directly used to analyze movements of 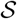 with respect to 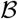 without knowledge of the translational and rotational kinematics of 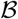 with respect to 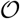.

#### Rotational motion

The angular velocity of 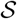 with respect to 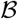 is given by:

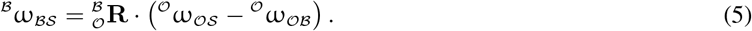

Combining Eqs. (5) and (4), we can obtain 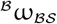 from the IMU’s gyroscope signal 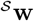

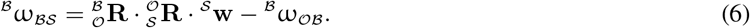

Thus, similar to the translational case, when 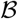 is not fixed, and one needs to isolate the kinematics of a specific body part, the kinematics of the reference frame 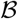 must be accounted for.

#### The special case where the body reference frame is fixed

In many experimental setups in movement neuroscience and neurorehabilitation, 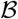 is fixed with respect to 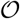 (e.g., trunk movements are restricted). This ensures that the position and differential movement kinematics of the body parts of interest 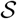 with respect to 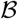 are related to the kinematics with respect to 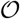 through fixed affine and linear transformations, respectively. If 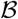 is fixed with respect to 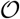 (i.e. 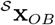 and 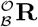 are constant), 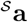 reduces to

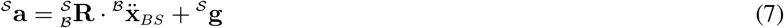

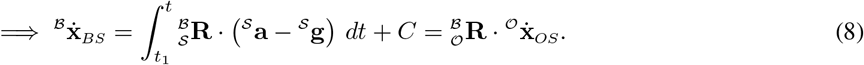

We will only discuss discrete movements in our analysis, i.e. movements that are bookended by postures where all derivatives of position are zero, which implies that *C* = **0**.

Then, the *movement smoothness* of the translational motion of 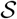, when 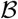 is fixed, is

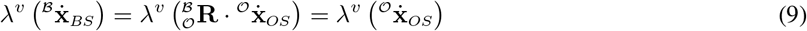

for both SPARC and LDLJ-V, since 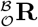 is fixed with respect to time.

For rotations, similar to the translational motion case, if 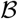 is fixed, i.e., 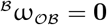, then,

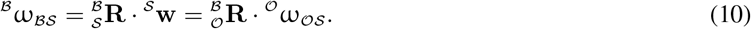

Assuming that the smoothness of rotational movements can be analyzed in the same way as translational movements, Eq. (10) allows us to estimate movement smoothness, when 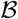 is fixed, as

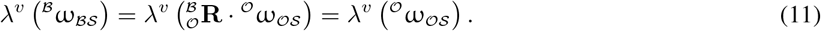

## 3 Movement smoothness from IMUs: challenges

The most straightforward approach to estimate movement smoothness from IMU data would be to reconstruct translational and angular velocities with respect to 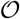, i.e. 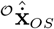 and 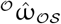, from the IMU measurements (Fig. 3; Eqs. (3) and (4)). Then, the smoothness could be computed using SPARC or LDLJ-V from these reconstructed kinematic variables as expressed in Eqs. (9) and (11). However, in practice, errors in attitude reconstruction of *S* with respect to 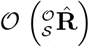 can result in severe errors in movement smoothness computed by SPARC and LDLJ-V. In the following, we analyze these issues for translational and rotational movements, their effects on the SPARC and LDLJ-V measures, and propose alternative ways to compute movement smoothness from IMU data. We will only consider discrete movements in our analysis, i.e. movements that start and end with a static posture (Hogan and Sternad, 2007). For such movements, the velocity is zero at the start and end of the movement

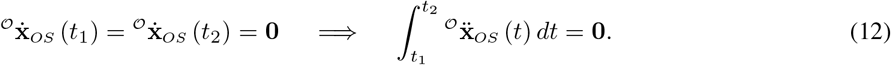

**Figure 3:**
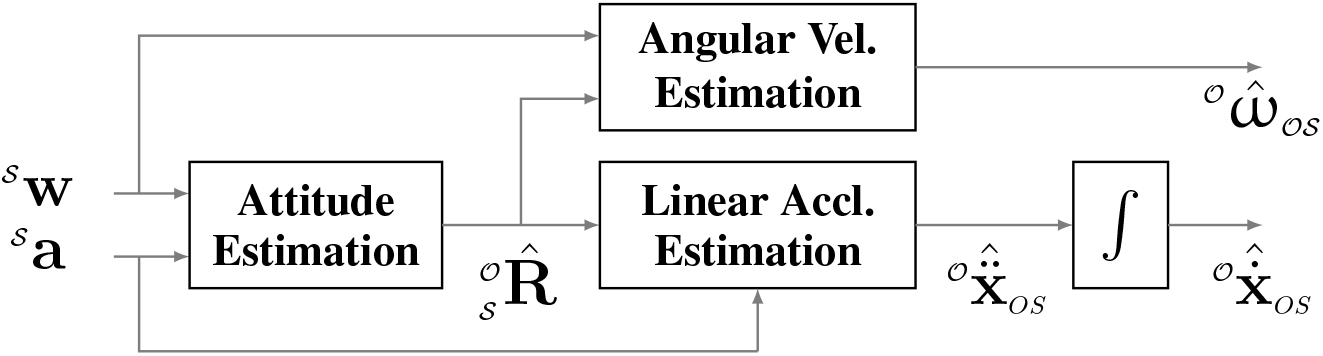
Reconstruction of kinematic variables of interest for estimating smoothness from IMU data.

### 3.1 Translational motion

To analyze the effects of kinematic reconstruction errors, the estimated linear acceleration from IMU data can be expressed as:

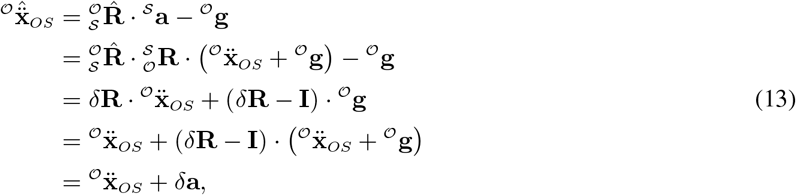

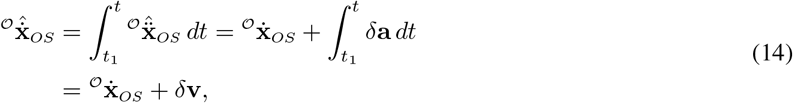

where 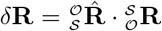 is the error in orientation reconstruction, and *δ***a**, *δ***v** are the linear acceleration and velocity reconstruction errors, respectively.

Thus, in the case of translational motion, using the SPARC 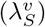 or LDLJ-V 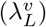 is problematic because of drift in 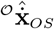 caused by integration of *δ***a** and noise in the sensor data (which is not considered in these equations). In movement analysis, the magnitude of gravity 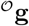 is much larger than linear accelerations of typical human arm movements, which can result in significant drift in 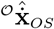 due to reconstruction errors; this problem is exacerbated in the case of slow, long-duration movements.

#### Proposed solution: LDLJ-A for acceleration data

As previously mentioned, applying SPARC or LDLJ-V to acceleration signals is semantically incorrect to obtain the smoothness of the movement represented by these acceleration signals. It is not clear how one would modify SPARC to work with acceleration data, given that there is no simple relationship between the magnitudes of velocity and acceleration. However, we can obtain jerk from acceleration, and choose an appropriate scaling factor to allow this measure to work directly with acceleration data, which results in the following new measure (LDLJ-A) applied to acceleration (see Appendix B for details about its properties).

##### LDLJ (based on acceleration; LDLJ-A)

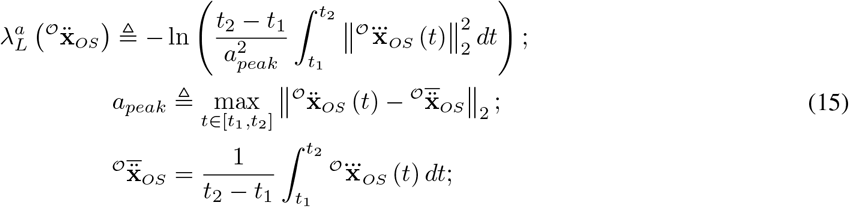

where 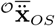 is the mean acceleration, which is zero for discrete movements. However, the mean of accelerometer data will not be zero, in general, even for discrete movements due to gravity and data segmentation practices to determine movement onset and termination. Removing the mean in this case will ensure *a_peak_* is not overestimated. It should be noted that the mean value will not affect the jerk integral and thus it need not be removed.

### 3.2 Rotational motion

Dealing with rotational motion from IMU data is easier, since: (a) the sensor directly measures angular velocity via the gyroscope, albeit in the local sensor reference frame, and (b) the signal is not affected by gravity.

For SPARC, the sensor orientation is irrelevant, since the operator ||·||_2_ is rotation invariant. The movement speed can then be computed directly from the gyroscope data, i.e., 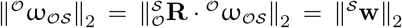. Therefore, 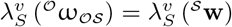.

In the case of LDLJ-V, 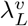 requires 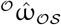, which is affected by the attitude reconstruction error as:

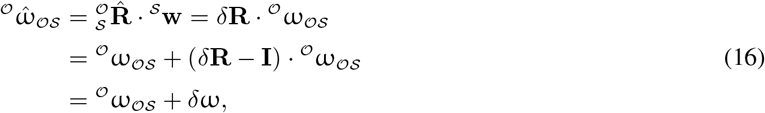

where *δ*ω is the rotational velocity reconstruction error.

Since the rotational jerk is estimated by double differentiation 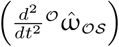, it is important to understand how errors in the orientation estimation affect the computation of LDLJ-V from the gyroscope data.

We note that there is an alternative formulation to compute the magnitude of jerk (and thus LDLJ-V) that is unaffected by the sensor rotation (see Appendix D for details). Thus, in principle, LDLJ-V can also be applied to gyroscope data to precisely compute smoothness. However, this formulation was found to be sensitive to practical implementations (e.g. choice of numerical differentiation methods, sampling frequency) and thus, needs further investigation before general recommendations can be made for its use in smoothness analysis.

## 4 Validation

As detailed in the previous section, when working with reconstructed acceleration data 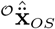, there will be errors in the jerk and *a_peak_* computation in 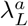 due to the acceleration reconstruction error *δ***a** (Eq. 13). Similarly, applying the LDLJ-V on rotational velocity measured from a gyroscope will result in errors in jerk computation (due to *δ*ω), which will affect the smoothness estimate. Thus, from a practical point of view, it is important to understand the effect of these reconstruction errors on the smoothness estimates of translational and rotational movements, to allow proper interpretation of the results from smoothness analysis using reconstructed data. In this work, we will only focus on applying LDLJ-A on translational movements, and the SPARC and LDLJ-V on rotational movements. This section first presents the results from the validation through simulated data, followed by the experimental validation.

### 4.1 Validation using simulated data

To test the proposed methods, we simulated discrete point-to-point movements, which were generated as 1*s* minimum jerk trajectories with varying number of via-points. We varied the following parameters:

i. **Number of via-points**: change in the number of via-points (spatio-temporal constraints) can lead to changes in movement smoothness. We used a subset of simulated discrete point-to-point movements from the analysis presented in Appendix B; twenty-five different trials with different via-points (*N_via_* = {1, 2, 5, 10}) were selected for this analysis. The distance between the start and end positions was set to 15*cm*.
ii. **Movement duration**: for a fixed movement amplitude, movement duration controls the magnitude of the accelerations and velocities. The accelerometer data from shorter duration movements will be dominated by the linear acceleration component 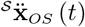, while longer duration movements will be dominated by the gravity component 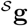. Four different movement durations *T* = {2.5, 5, 10, 20} *s* were simulated for each movement trial by time-scaling the simulated movements of 1s duration. The velocity and acceleration of a movement of duration *Ts* were derived as

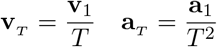

where **v**_1_ and **a**_1_ are the velocity and acceleration of the 1*s* duration movement.
iii. **Attitude reconstruction error**: simulating different attitude reconstruction errors can be done by generating a stochastic time-series of rotation matrices. This was done through the following parametrization

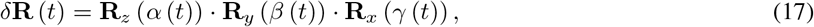

where **R**_*x*_ (·), **R**_*y*_ (·), and **R**_*z*_ (·) are the elementary rotation matrices about the *x*, *y*, and *z* axes, respectively; *α* (*t*) *, β* (*t*), and *γ* (*t*) are Euler angles that determine the amount of rotation about each axis. The time-series of these Euler angles were realizations of Gaussian Brownian noise (integration of white noise), which were low-pass filtered through a moving average filter (window size of 0.5*s*). These angles were scaled such that max_*t*_ |*α* (*t*)*|* = max_*t*_ |*β* (*t*)*|* = max_*t*_ |*γ* (*t*)*|* = *θ_max_*; *θ_max_* determines the maximum amount of rotation error represented by *δ***R**. Three different values were used for *θ_max_* ∈ {5°, 25°, 50°}. For each movement trial with a fixed number of via points and duration, we simulated five different realizations of the Euler angles for each level of *θ_max_*.

The simulated movements were used to generate reconstructed accelerometer and gyroscope data as

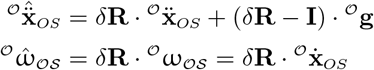

where 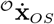 and 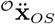 are from the simulated movements, and *δ***R** is from the simulated sensor orientation reconstruction error (Eq. 17). For the gyroscope data, we simply treated the simulated movements as rotational movements, and used the velocity associated with these movements 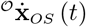 as the angular velocity 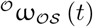.

The summary of the values for the different factors used in the simulation analysis are presented in Table 1, which resulted in a total of 6000 movements. Movement smoothness of the simulated reconstructed movements of the accelerometer and the gyroscope were estimated by:

- Applying 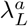 to the original movement acceleration 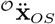 and the reconstructed acceleration 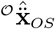.
- Applying 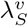 and 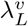 to the original angular velocity 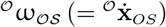 and reconstructed angular velocity 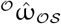.

**Table 1:**
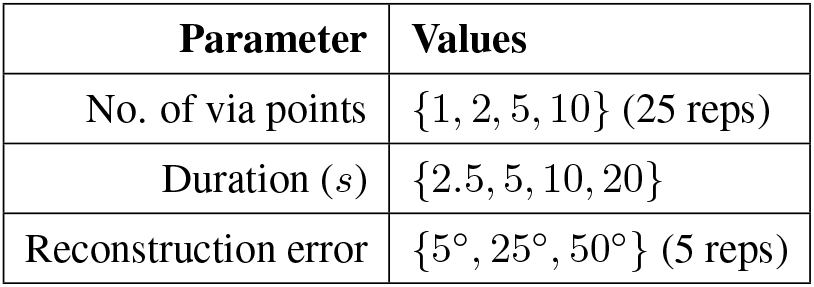
Summary of parameters used to analyze the effect of reconstruction error on movement smoothness. A total of 6000 simulated movements were generated.

| Parameter | Values |
| --- | --- |
| No. of via points | $\{1, 2, 5, 10\}$ (25 reps) |
| Duration (s) | $\{2.5, 5, 10, 20\}$ |
| Reconstruction error | $\{5^\circ, 25^\circ, 50^\circ\}$ (5 reps) |

The relative error *ϵ* between the smoothness of the original movement *λ* (**y**) and the reconstructed movement 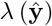 was calculated as:

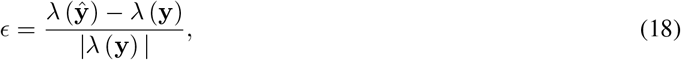

where *λ* (·) is the smoothness measure, **y** and 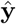 are the original and reconstructed movement kinematic variables of interest, respectively.

In the case of translational movements, the relative magnitude of the true linear acceleration with respect to gravity is useful to gauge the relevance of the sensor measurement to quantify the movement in question. A rough estimate that can be obtained from the accelerometer data is the sensor-to-gravity ratio (SGR):

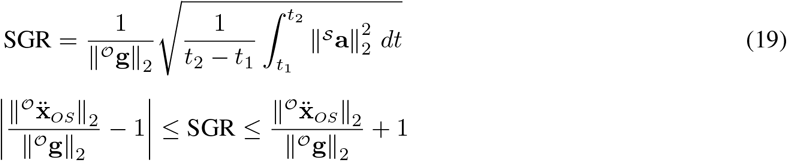

SGR can only assume non-negative real values, and provides an approximate idea about the relative contributions of the linear acceleration and gravity components. An accelerometer that is free-falling with respect to an earth-fixed reference frame will register **0** acceleration on the accelerometer, and thus have *SGR* = 0. Values of SGR farther from 1 indicate an increasing contribution from the movement’s linear acceleration.

The summary of the smoothness analysis of the simulated accelerometer data is shown in Fig. 4. The smoothness values of the simulated original and reconstructed movements were roughly related (Fig. 4(a)). The Pearson correlation coefficient was 0.713 for all 6000 movements, and 0.945 for the subset of movements with *SGR* ≥ 1.05. Thus, as expected, movements with larger linear acceleration have better smoothness estimates. Further, smoothness of reconstructed movements with SGR ≥ 1.05 were closer to the actual smoothness value (Fig. 4(b)). The highest error in this case was ≈ 40% off the actual value, whereas for movements with SGR below 1.05 it can be ≈ ±60% off the actual value.

**Figure 4:**
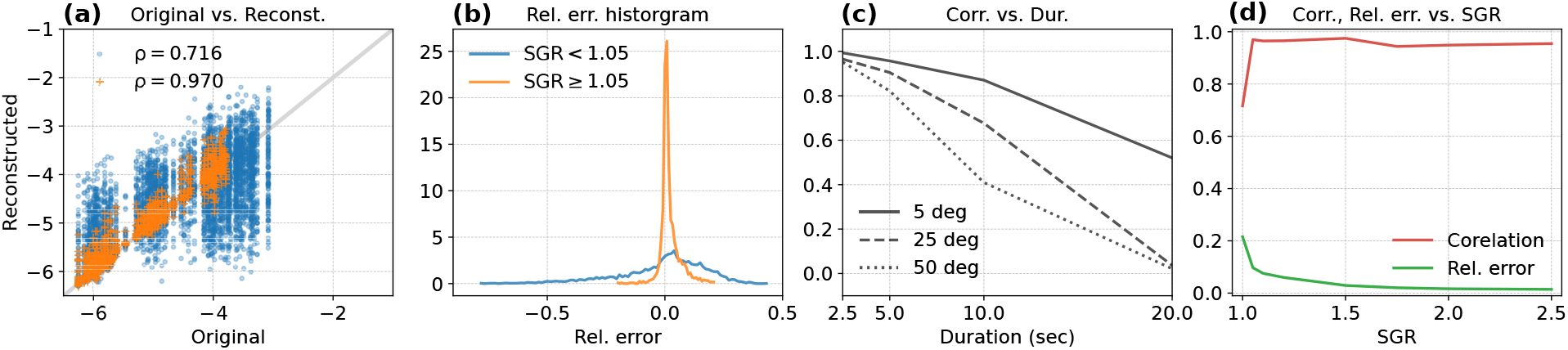
Evaluat on of 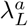 to estimate smoothness of translational movements: simulated original 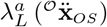 and reconstructed 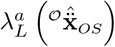 movement smoothness. (a) Smoothness of original versus reconstructed kinematics. The blue dots correspond to each of 6000 simulated movements, while the orange ‘+’ are movements with *SGR* > 1.05. The *ρ* values are their respective Pearson correlation coefficients. (b) Density histogram of the relative error between smoothness of the original and the reconstructed movements for *SGR* < 1.05 (blue curve) and *SGR* > 1.05 (orange curve). (c) Correlation between smoothness of original and reconstructed movements as a function of movement duration for different attitude reconstruction errors. (d) Correlation and relative error between smoothness from original and reconstructed movement for all movement above a particular SGR value along the *x*-axis.

Improved reconstruction resulted in better estimates (Fig. 4(c)). For all movement durations, the correlation coefficient increased with decreasing reconstruction errors. For a given level of reconstruction error, the correlation decreases with duration because of the decreasing contribution of the movement’s linear acceleration relative to gravity. This means that smoothness estimates are significantly off from the original smoothness values for slow movements that are poorly reconstructed.

Finally, we analyzed how the correlation and relative error between the smoothness of the original simulation and reconstructed movements varied as a function of SGR. This was done by first selecting all movements with SGR greater than or equal to a particular value, and then estimating the correlation coefficient and the relative error between 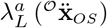 and 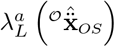 for this set of movements; these plots were generated for *SGR* ∈ [1.0, 1.05, 1.1, 1.2, 1.5, 1.75, 2.0, 2.5]. The relative error shows a monotonic decrease with increasing SGR. However, the correlation coefficient shows a non-monotonic increasing trend with increasing SGR. This is because of the limited number of movements with high SGR in the simulated data set, and the smaller range of smoothness values these movements assume. For example, movements with SGR ≥ 1.05 had original smoothness values mainly between −6.5 and −4, and this range decreased with increasing SGR.

For rotational movements (Fig. 5), as expected, there was perfect correlation between smoothness of the original and reconstructed movements for the SPARC (*ρ* = 1); however, the LDLJ-V has a much lower correlation of *ρ* = 0.51 (Fig. 5(a)). The error in smoothness is larger for smoother movements, reaching up to 100% relative error (Fig. 5(b)), primarily due to the error in jerk calculation from 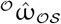.

**Figure 5:**
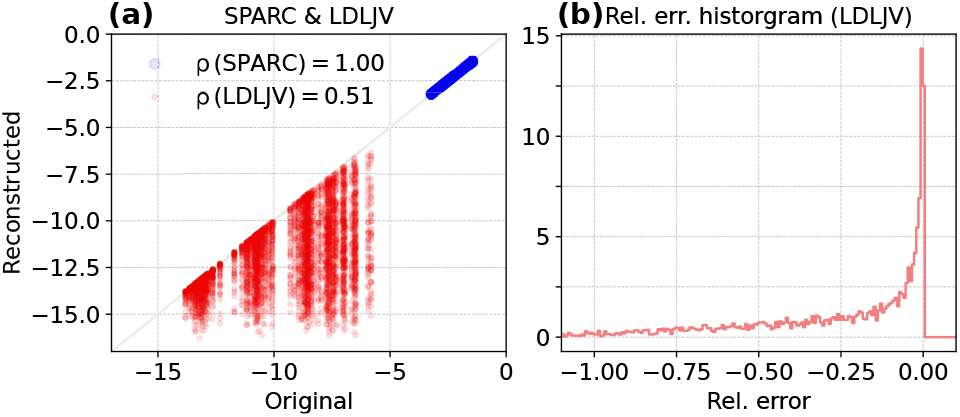
Evaluation of 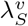 and 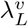 to estimate smoothness of simulated rotational movements. (a) Smoothness of original 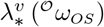 versus reconstructed movements 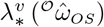; the red dots and blue circles correspond 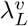, and 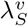, respectively. (b) Density histogram of the relative error between smoothness of the original and the reconstructed movements 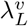.

### 4.2 Validation using human movement data

To evaluate the consistency of estimating movement smoothness in practice, we applied the SPARC, LDLJ-V and LDLJ-A to human arm movement data collected simultaneously with an optical passive marker-based motion capture system (MoCap) and IMUs. We used a set of 250 upper-limb movements, collected from 4 post-stroke individuals with different levels of arm impairment. Data were not collected for the specific purpose of this paper; they were collected during the patients’ stay at the cereneo center for Neurology and Rehabilitation (Vitznau, Switzerland) as part of a different research study. We only used data from patients who authorized ‘further use of data’.

The kinematic dataset consisted of patients performing all tasks of the ARAT (Arm Research Action Test) with both their right and left arms. An optical motion capture system (Qualysis, Göteborg, Sweden) and one IMU worn on each wrist (ZurichMOVE, ETH Zurich, Switzerland) were used to track the kinematics of both arms. Tso track the movement simultaneously with the two systems, a cluster of passive-reflective markers was attached to each of the IMUs as shown in Fig. 2(b).

Marker data for each ARAT task were tracked and exported to Visual3D (C-Motion, Germantown MD, USA), then low-pass filtered with a zero-lag 2^*nd*^ order Butterworth filter with cut-off frequency at 20Hz. A rigid-body model was used to calculate the position 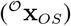, angular velocity 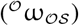 and orientation 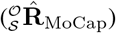 of each cluster of markers representing the IMUs.

Data from the IMUs were a collected as continuous data streams, and the MoCap data were used to align the data in time and segment it into each ARAT task. Data from each IMU consisted of acceleration 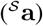, angular velocity 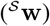, and quaternions representing the orientation of the IMU, which were used to compute the orientation of the sensor 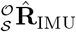.

Linear velocity was obtained by numerical differentiation of position data 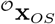 (MoCap data) or integration of the rotated accelerometer signal minus gravity 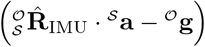 (IMU data)

Smoothness measures were computed for both IMU and MoCap data. For SPARC, the adaptive cut-off frequency was chosen to be less than or equal to 6Hz (i.e. *ω_c_* = 12*π* in Eq. (1)), and the signals were low-pass filtered with a zero-lag 2^*nd*^ order Butterworth filter with a cut-off frequency at 6Hz. To further understand the effect of errors in the rotational matrix on smoothness estimates, we generated simulated errors similarly to section 4.1 For each of the 250 upper-limb movements, 100 stochastic time-series rotation matrices (**R**_noise_) with varying Euler angles ranging from 5 deg to 50 deg were generated, resulting in 250,000 movements.

The agreement between IMU and MoCap smoothness values was quantified as a percentage relative to the smoothness quantified from MoCap (relative error), as MoCap is the ‘gold standard’ for movement analysis. Limits of agreement are reported as 1.96 times the standard deviation of the computed values and are shown in the figures with a 95% confidence interval region (Giavarina, 2015).

The summary of the results from this analysis is presented in Figs. 6 and 6. For linear velocity, both SPARC (Fig. 6(a)) and LDLJ-V (Fig. 6(b)) resulted in highly uncorrelated values between MoCap- and IMU-based estimates (SPARC, *ρ* = −0.11; LDLJ-V, *ρ* = −0.10). As expected (see section 3.1), the drift caused by integration significantly affected the smoothness estimated from the IMU data. For SPARC, the drift causes a relative increase in the DC frequency component of the speed signal. This, in turn, decreases the adaptive cut-off frequency *ω_c_* (eq. 1), resulting in a shorter spectral arc length, and thus a smoother estimate. For LDLJ-V, the drift results in the over-estimation of *v_peak_*, resulting in a smaller normalization factor, and thus a smoother estimate.

**Figure 6:**
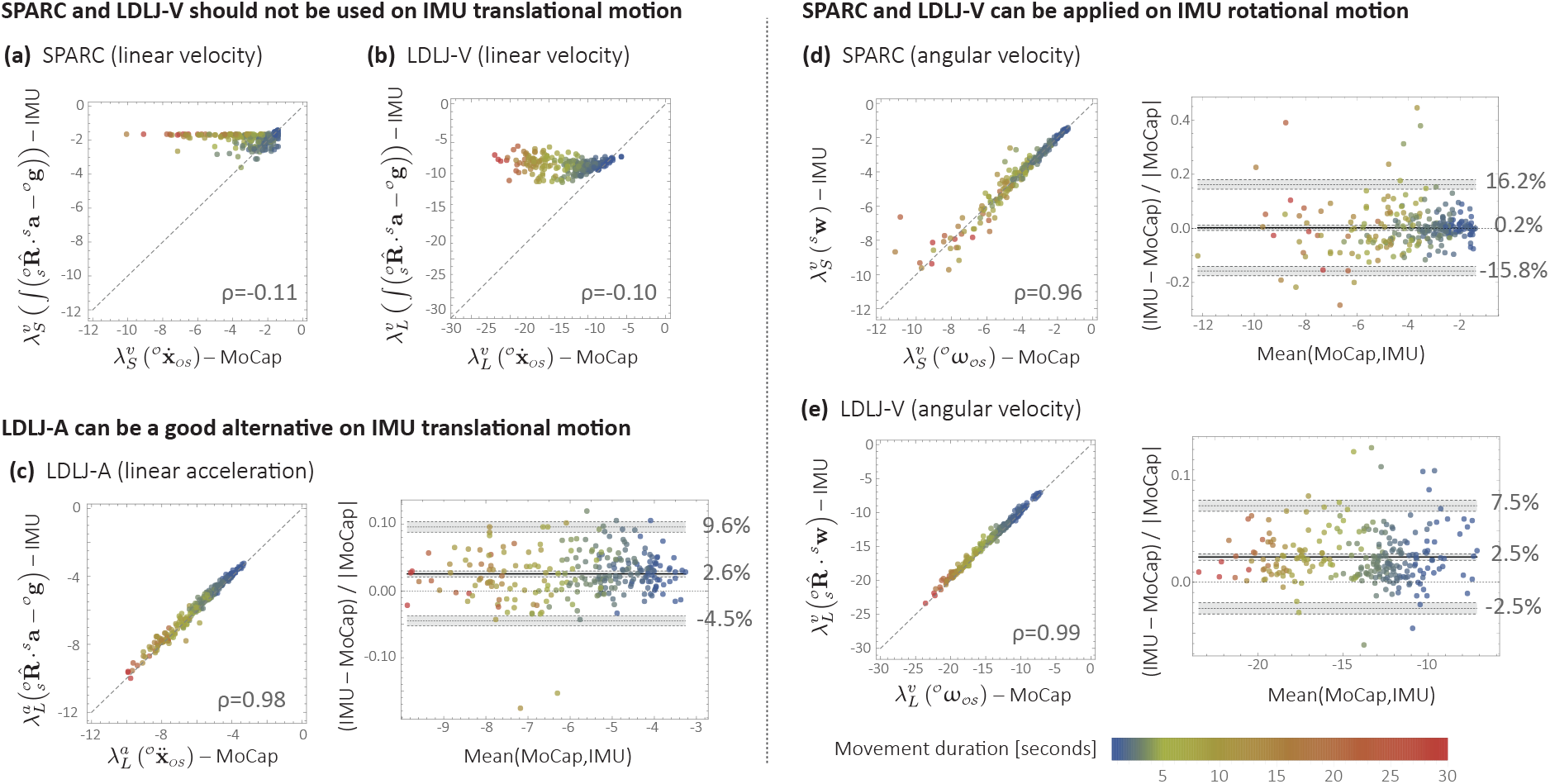
Comparison between smoothness estimates from Motion Capture (MoCap) and IMU data during upper-limb movements of stroke survivors.

MoCap- and IMU-derived smoothness computed using LDLJ-A (Fig.6(c)) were highly correlated (*ρ* = 0.98).The limits of agreement indicate that IMU-derived estimates were smoother than those derived from MoCap, with a mean bias of 2.6% (range from −4.5% to 9.6%). One explanation for the bias could be the amplification of noise in the MoCap position data due to triple numerical differentiation needed for computing LDLJ-A, resulting in smaller smoothness values. The overestimation of smoothness computed from IMU data was also seen with simulated movements in Fig. 4(a), where movements with smoothness values less than −4 are mostly on or above the identity line; the smoothness values of all movements in Fig. 6(c) are also lower than −4. Another possible explanation could be the over-estimation of *a_peak_* because of imperfect removal of gravity.

The reduced spread of points in Fig.6(c) compared to Fig. 4(a) and the high correlation observed were obtained even though a large number of experimental movements had an *SGR* < 1.05. We believe this was likely due to small reconstruction errors in 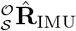 that could be obtained from the this particular IMU brand. Indeed, when simulated reconstruction error was added to 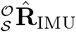 (see Fig. 7(a)), the relative error between MoCap and IMU increased significantly overall.

**Figure 7:**
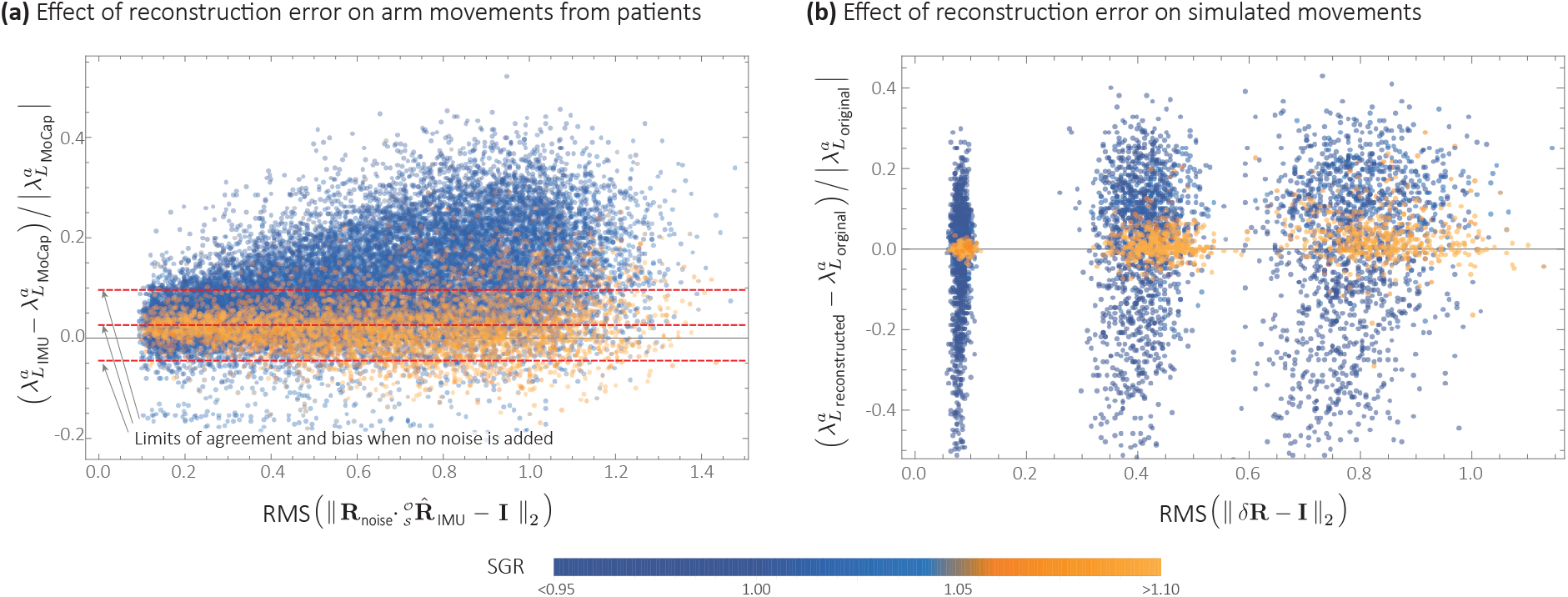
Effect of orientation reconstruction error on LDLJ-A. (a) Agreement between MoCap and IMU from experimental data, when simulated noise is added to the rotation matrix 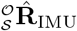 obtained from patient data. (b) Relative error *ϵ* (defined in Eq. 18) when simulated reconstruction error *δ***R** is added to synthetic movement data. Here, the metric *RMS* (||*δ***R** − **I**||_2_) is used as measure of orientation reconstruction error.

Fig. 6(d) and Fig. 6(e) show SPARC and LDLJ-V computed on the angular velocity from MoCap 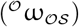 and the reconstructed angular velocity from the IMU’s gyroscope 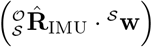. The MoCap- and IMU-derived smoothness values were highly correlated for both SPARC (*ρ* = 0.96) and LDLJ-V (*ρ* = 0.99). While SPARC resulted in no significant bias in relative error (0.2%), LDLJ-V had a relative error bias of 2.5%. On the other hand, the limits of agreement were smaller for LDLJ-V (−2.5% to 7.5%) compared to SPARC (−15.8% to 16.2%). The fact that SPARC resulted in poorer levels of agreement than LDLJ-V was surprising, given that: (a) with angular velocity, SPARC is unaffected by rotation 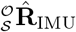; and (b) SPARC has been shown to be more robust to noise than LDLJ-V (Balasubramanian et al., 2012, 2015).

## 5 Discussion

Quantification of *movement smoothness* is considered to be of great importance in movement sciences and biomechanics, with smoother movements implying unimpaired, skilled behavior, and its change is associated with changes in motor control. Thus, the goal of smoothness analysis is to gain insights about the underlying mechanisms that generated the observed motor behavior. Several groups have proposed that the construct of *movement smoothness* can be understood through the concept of submovements: *“…smooth movements are movements composed of a few submovements that are closely spaced in time, while unsmooth movements would result from the superposition of a larger number of submovements with loose temporal packing”* (Balasubramanian et al., 2012). Therefore, a consistent movement smoothness measure should have a *“…monotonic response to the motion characteristics so that the smoothness measure decreases with the number of submovements and the inter-submovement interval.”* (Balasubramanian et al., 2012). Using this as an operational definition, the current recommended smoothness measures - SPARC and LDLJ-V - have been developed and validated using velocity kinematics as the representation of the movements of interest. This paper shows that the SPARC and LDLJ-V cannot be arbitrarily used directly with other movement variables (e.g. position, acceleration, etc.). The numbers resulting from simply replacing the velocity term by any other movement variable (or any signal) cannot be interpreted as *movement smoothness*. For example, applying the SPARC on an arbitrary signal *s* (*t*) will produce a number that measures the level of ‘undulations’ in that signal. This number can be interpreted as smoothness of a movement *M* only if *s* (*t*) represents the speed profile of the movement *M*. This issue is highly relevant for movement data for which velocity cannot be easily estimated. The data from an IMU is a typical example.

IMUs provide an alternative mode of sensing movement kinematics that is cheaper and more convenient than ‘traditional’ methods, such as motion capture systems. However, in contrast to these systems, IMUs do not directly measure position kinematics, have a time-varying frame of reference, and are noisy – which make deriving high-quality velocity and position kinematics non-trivial. These issues put into question the appropriateness of IMU data to quantify *movement smoothness* in practice.

In this paper, the difficulties of using IMUs for movement smoothness analysis on translational and rotational motions was presented through a systematic analysis. The ultimate goal of this work was to develop recommendations that can help standardize methods for estimating *movement smoothness* using not only IMUs, but across different movement measurement technologies. A summary of recommendations for the measures to use for analysing movement smoothness using IMU data is in Fig. 8.

**Figure 8:**
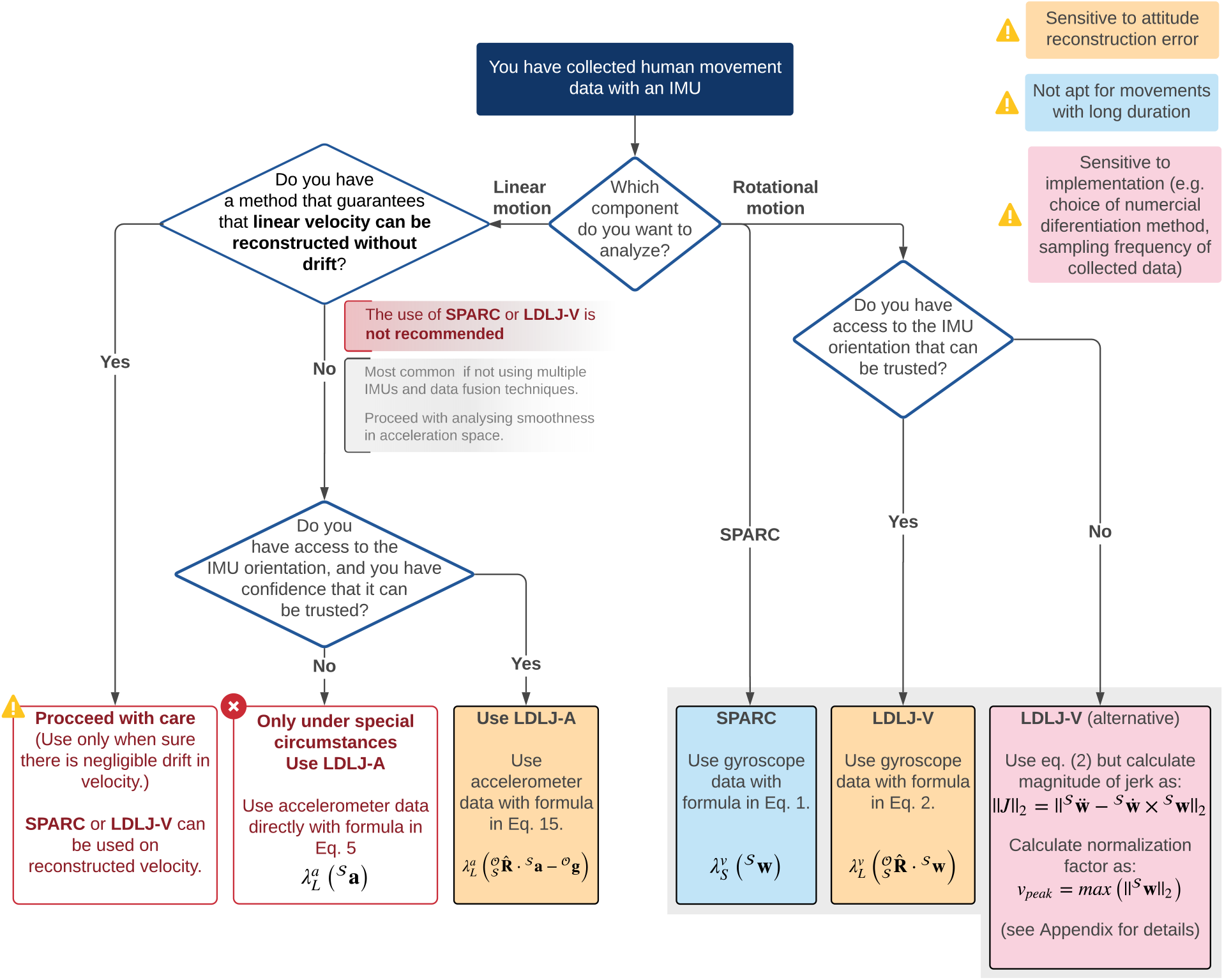
Summary diagram of recommendations for analysing movement smoothness of discrete movements (i.e., with a clear start and end with zero velocity) using IMU data.

### 5.1 Translational movement smoothness

Estimation of linear velocity is required to compute SPARC and LDLJ-V. This computation is complicated because it requires integration of the accelerometer signals, which are affected by rotational movements of the sensor and are noisy. Even if one has a good estimate of the IMU’s rotation, reconstruction errors and accelerometer noise can translate into drift in the estimated linear velocity. This, in turn, affects the reliability of the smoothness measures (see Fig. 6(a-b)). Thus, **SPARC and LDLJ-V should not be used on translational velocity kinematics obtained from an IMU**. There are several techniques that can be used to obtain improved estimates of translational kinematics from fused information from multiple IMUs (Kok et al., 2017), which could enable the use of SPARC and LDLJ-V on IMU-based data. However, such pose estimation methods rely on application-specific models, constraints, and assumptions, which can introduce systematic bias in the velocity estimate of the underlying motion. Thus, depending on the selected approach, the same motion, processed by different algorithms, can result in significantly different movement smoothness estimates. To our knowledge, there have been no studies evaluating the sensitivity of smoothness measures on such translational velocity reconstruction algorithms from IMUs. Thus, at this point, estimating smoothness with SPARC and LDLJ-V using such algorithms are best avoided until their reliability is established.

*Movement smoothness* is related to the concept of intermittency of the executed motion, which is also associated with the idea of temporal dispersion of submovements. One interpretation of movement intermittency is the presence of *movement arrest period* (MAP), which is a continuous interval of time where there is no movement, i.e., where all derivatives of position are uniformly zero. **Movement velocity is the most direct indicator of movement intermittency caused by MAPs**. This is because all time intervals with uniformly zero velocity are MAPs, while non-zero velocities indicate movement. This unique relationship, however, is lost with higher derivatives (acceleration, jerk, etc.), e.g., intervals with zero acceleration can either indicate an MAP or an interval with constant velocity. The strong reliance of SPARC on MAPs is the main reason why SPARC applied to acceleration data looses its interpretability as a movement smoothness measure (see Appendix C).

On the other hand, the jerk-based measure does not directly rely on MAPs to quantify intermittency, but rather does this through the jerk term, i.e., changes to the movement acceleration. Since jerk can be derived from acceleration data, jerk-based measures can potentially be calculated from acceleration with the appropriate modifications in the scaling factor. This resulted in the LDLJ-A proposed in Eq. 15, which can be used with acceleration data.

The analyses presented in this study show that the **LDLJ-A can be a good alternative to estimate translational movement smoothness from IMU acceleration data**. However, its use requires a reasonably well reconstructed movement acceleration signal, which is determined by the magnitude of error in the IMU’s attitude reconstruction, and the relative magnitude of the movement’s translational acceleration with respect to gravity. This was illustrated with simulated (Fig. 4(c) and Fig. 4(d)), and experimental data (Fig. 7). Unfortunately, the extent of the reconstruction error and the relative magnitude of linear acceleration with respect to gravity are often unknown. In such a scenario, the SGR metric (Eq. 19) can be used to provide some confidence in the smoothness estimates. Movements with an SGR of at least 1.05 were found to be less sensitive to different amounts of reconstruction errors (Fig.4(a) and Fig.7).

Estimating movement smoothness from an accelerometer without an estimate of its orientation should ideally be avoided except under the special circumstance where there is a good reason to believe that the sensor did not undergo much rotation, and the SGR is greater than 1.05 (the larger the SGR, the better the estimates). In this very specific case, one could potentially apply the LDLJ-A 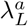 measure directly on the accelerometer data 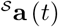.

### 5.2 Rotational movement smoothness

Analyzing rotational movements from IMU data is simpler than the translational case because gyroscopes directly measure rotational velocity and are unaffected by gravity. **SPARC can be applied to gyroscope data without any modifications**. LDLJ-V needs the gyroscope data to be corrected for sensor rotation (Eq. 16), and thus, can be affected by reconstruction error. We note that there is another approach that can theoretically circumvent the need for orientation correction (described in Appendix D), but this approach suffers from some practical implementation issues which need further investigation.

In experimental data, we found that LDLJ-V computed from IMU and MoCap had better agreement than SPARC. This is in contrast to our previous studies (Balasubramanian et al., 2012, 2015), in which we consistently found that SPARC has better reliability than LDLJ-V, especially when it comes to noisy data. When looking closer at the SPARC results, we observed that the performance was poorer for less smooth (and also longer duration) movements. Movements that exhibited poorer levels of agreement had slight differences between the IMU and MoCap magnitude spectrum curves (at frequencies lower than 6Hz), which, when integrated, resulted in significant differences in spectral arc length. However, LDLJ-V was insensitive to those differences because of the logarithmic transformation which compressed large differences in the dimensionless jerk (the argument of the log function in Eq. 15) between the IMU- and MoCap-based estimates. This problem is observed primarily for movements with a large number of submovements or undulations; such complex movements were not considered in our previous studies, and thus, this characteristic of SPARC was unnoticed. Future work could investigate possible transformations on SPARC (similar to the logarithm) that can alleviate this behavior.

One must not forget, that the good and bad results for LDLJ-V and SPARC reported from experimental data only reflect the characteristics of the specific dataset presented here. It remains to be seen how these measures behave on a different set of movements. Since the correlation between the IMU- and MoCap-derived smoothness using both SPARC and LDLJ-V were very high, we cannot recommend the use of one measure over the other - both approaches seem good candidates for analysing movement smoothness of rotational movements measured by IMUs.

### 5.3 Limitations

It is crucial to bring to light some of the shortcomings of the current study to ensure the results are interpreted appropriately.

1. The work presented is only an initial step towards standardizing methods for estimating **movement smoothness** using IMUs that is consistent with other movement measurement technologies. The preliminary validation of the different proposed approaches in this work are based on a small set of simulated and experimental data. Thus, it is possible that part of the study outcomes are specific to these datasets. The simulated and experimental movements had a wide range of smoothness values indicating that a broad range of behaviors were included. Nevertheless, the recommendations made in this study must be validated using a larger dataset with more subjects and a wider range of movement tasks.
2. The tracking of rotational movements with MoCap was found to be quite noisy, which could have been the reason for the bias and the wider limits of agreement for the different measures. A future validation study must quantify the noise characteristics of the IMU and the MoCap systems to better evaluate the performance of the different smoothness measures.
3. While evaluating the effect of reconstruction error *δ***R**(*t*) on smoothness estimates, we only considered the maximum reconstruction error, measured in terms of Euler angles of *δ***R** or the RMS value of induced 2-norm of the matrix *δ***R** − **I**(Fig. 7(b)). However, it must be noted that the smoothness of the reconstructed movement is not only affected by the magnitude of reconstruction error, but also the exact time profile of reconstruction error (i.e., ||*δ***R**(*t*) − **I**||_2_) and its relationship to the actual movement. Thus, it is possible that results shown in Fig. 4 would be different if we had used a different approach for generating the Euler angles. However, it can be shown that the effect of the time profile of ||*δ***R**(*t*) − **I**||_2_ will reduce as the amount of reconstruction error diminishes.

## 6 Conclusion

In this paper, we carried out a systematic analysis of applying different smoothness measures to discrete movement data measured by an IMU, with the aim of identifying the most appropriate methods for analysing smoothness with this type of data. Our results suggest that there appears to be no single optimal approach for analysing both translational and rotational movements measured by an IMU. The most appropriate method for estimating movement smoothness depends on two factors: (a) type of movement (rotational or translational), and (b) quality of the estimate of the IMU’s rotation. Based on our analysis, the recommendations for analyzing translational and rotational motions for smoothness analysis are summarized in Fig. 8. Future studies must evaluate these recommendations on larger dataset consisting of different types of movements.

## 7 Nomenclature

*r* scalar (lowercase italic letter)

**r** vector (lowercase bold letter)

**R** matrix (uppercase bold letter)

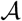 coordinate frame (calligraphy uppercase letter)

*A* origin of coordinate frame 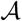

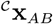 position vector of point *B* w.r.t point *A* expressed in frame 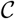

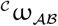 angular velocity vector of frame 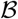 w.r.t frame 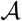 expressed in frame 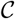

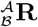 matrix representing a passive rotation that maps vectors expressed in frame 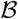 to frame 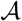

*t* time

*t*_1_ start time of a discrete movement (onset)

*t*_2_ stop time of a discrete movement (termination)

*g* gravitational acceleration constant (9.81*m/s*^2^)

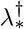 smoothness measure using method ∗ ∈ {*S, L*}, where *S* and *L* correspond to SPARC and LDLJ respectively, and evaluated from data type † ∈ {*p, v, a, j*}, where *p*, *v*, *a*, and *j* correspond to position, velocity, acceleration and jerk respectively

## Author Contributions

All authors contributed to the conception of the ideas and the theoretical aspects of the work. CS and AMC worked on the experimental setup and data collection used for validating the various smoothness measures. AMC analyzed the experimental data. SB worked on the simulation analysis of the different smoothness measures. All authors contributed equally to the interpretation of results, writing and reviewing of the manuscript.

## Acknowledgements

We thank Nathanael Jarrasse and Domenico Campolo for the fruitful discussions on the topic; and the cereneo Center for Neurology and Rehabilitation and Andreas Luft for support; thanks to Mario Widmer, Marianne Schesny, Alexandra Widmer and Meret Vogel for their help in data collection and marker tracking.

## Data Availability Statement

The code used for the generation and analysis of the simulated data in this paper is available at https://github.com/siva82kb/smoothness_from_imu.

## Appendix A Translational kinematics of a moving segment relative to another

The linear velocity of 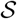 relative to 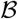 (Fig. 2) is given by:

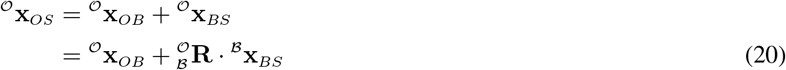

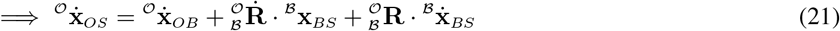

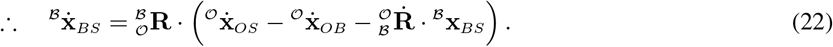

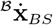 can be used to estimate the smoothness of the translational movements of 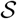 with respect to 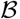. This is not the same as using 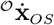, as it additionally contains the translational 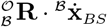 and rotational 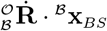 movement components of 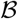. Thus, in general, 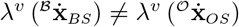, and they are equal only when 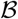 does not undergo any translational or rotational movements with respect to 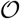, i.e. 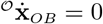 and 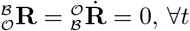.

We can easily derive 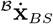 from position kinematics measured using an optical motion capture system. This, however, is not straightforward with IMU data. The linear acceleration of 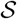 with respect to 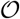 is given by deriving eq. (23):

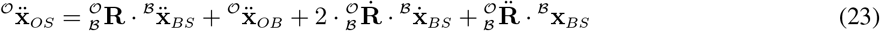

Combining eqs. 3 and 23, we get the accelerometer signal 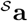,

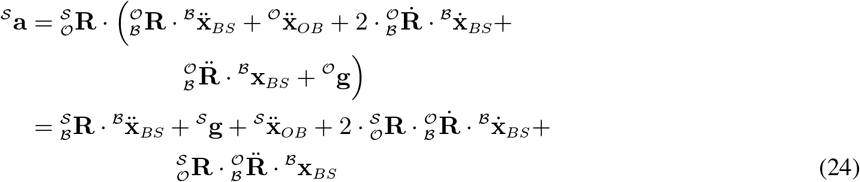

## Appendix B Analysis of the properties of LDLJ-A

The formulation of LDLJ derived directly from an acceleration signal is given by:

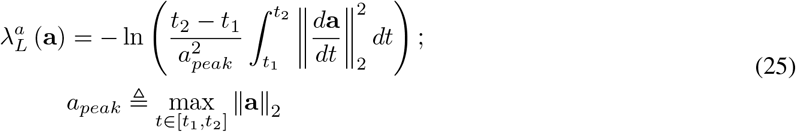

Comparing this formulation with 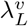 (Eq. 2), one can see that 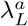 and 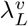 are different only in terms of the scaling factor 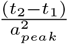 and 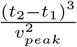, respectively, since

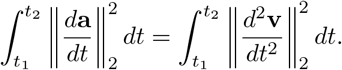

Thus,

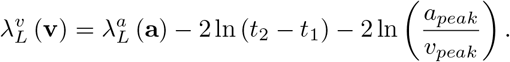

This relationship is non-linear due to the 2 ln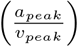 factor.

### Consistency of LDLJ-A

The consistency of the LDLJ-V and LDLJ-A measures were evaluated using simulated 1D movements, which were generated as a sum of submovements with varying inter-submovement intervals. The movement velocity and acceleration profiles were of the following form

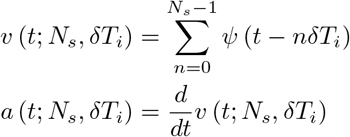

where *v* (·) and *a* (·) are the velocity and acceleration of the movement, respectively, 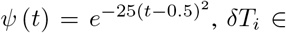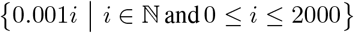 and *N_s_* ∈ {2, 4, 8}. The smoothness values of this set of movements was evaluated using the LDLJ-V 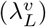 and LDLJ-A 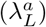 applied to the velocity and acceleration data respectively.

### Comparison of smoothness values 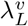 and 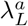

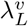 and 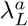 have roughly similar responses to changes in *δT_i_*, and show approximately monotonic responses to the change in submovement characteristics (Fig. 9). In general, for the same movement, the exact smoothness value computed from 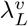 and 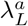 will not be the same. However, if there is an isomorphism between 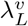 and 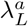, then both these measures would report the same relative smoothness between any two movements. For example, consider two movements *M*_1_ and *M*_2_ with velocity profiles **v**_1_ and **v**_2_, respectively. If there is a isomorphism between 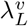 and 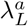, then

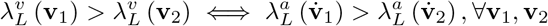

**Figure 9:**
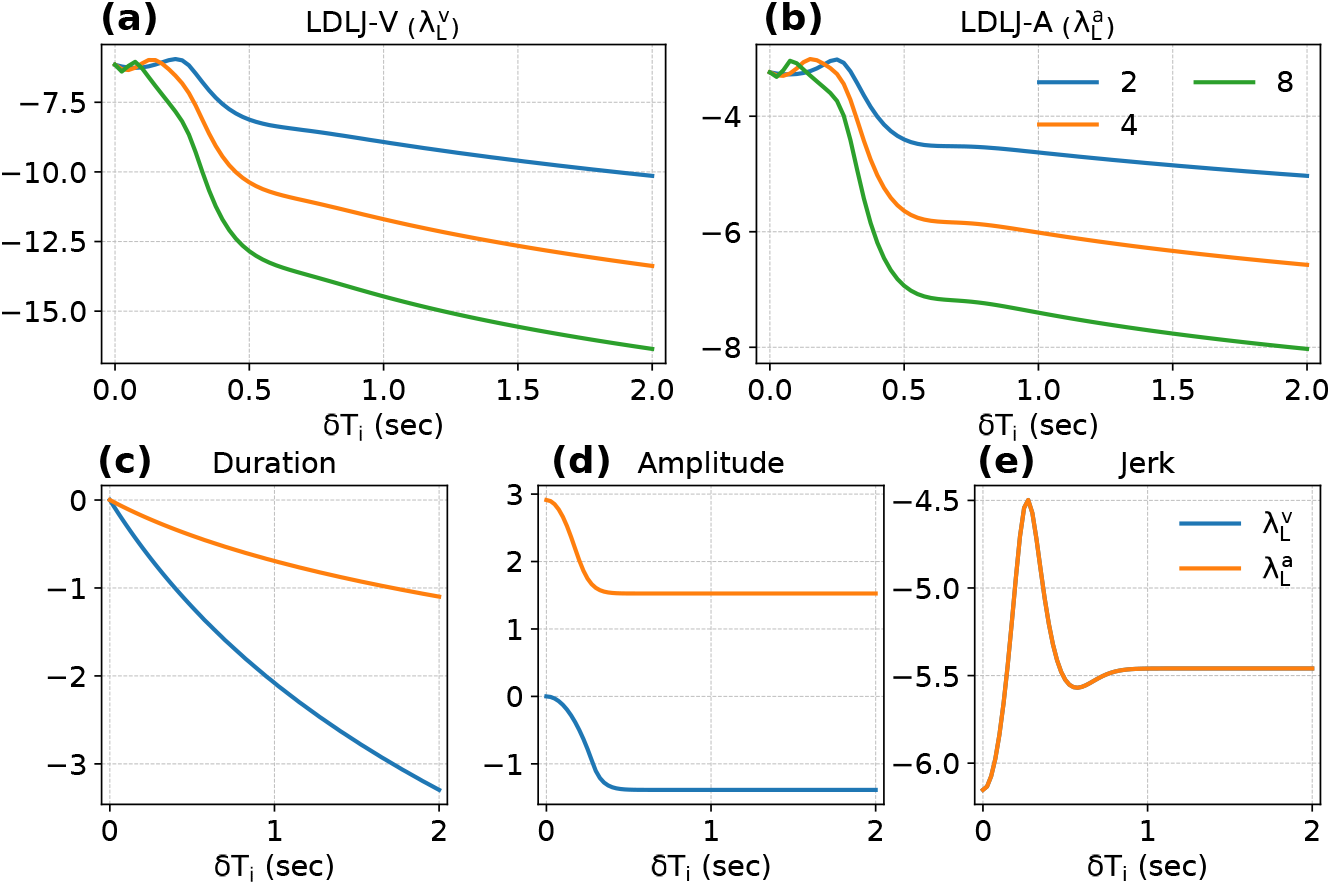
Consistency of 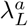. Subplots (a) and (b) depict the responses of 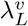 and 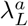, respectively, as a function of inter-submovement interval 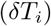. The response of the three individual terms (c) Duration (*T*), (d) Amplitude (*A*), and (e) Jerk (*J*) as a function of 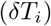 for 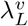 and 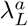.

To check if such an isomorphism exists between 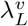 and 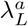, we simulated a set of discrete, minimum jerk, 3D movements through via-points. The problem was posed as a constrained optimization problem and was solved using the CVXPY solver in Python, similar to the approach used in Yazdani et al. (2012):

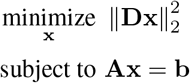

where **x** is a stacked vector of positions corresponding to the movement trajectory, **D** is the matrix performing the triple derivative to obtain movement jerk from **x**, and **A** and **b** represent the spatio-temporal constraints corresponding to initial, final, and via-points, and the zero initial, and final velocity and acceleration constraints. The solution **x**_*m*_ obtained using this procedure would possess the minimum squared jerk 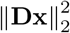 for the given spatial and temporal constraints.

The initial **x**_*i*_ and final **x**_*f*_ of all the movements were fixed to [0 0 0]^T^ and [0 1 0]^T^, respectively, and the movement duration was fixed to *T* = 1*s*. A set of *N* via-point spatio-temporal constraints 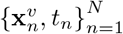 were randomly generated; 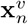 represents the position of the *n^th^* via-point and *t_n_* represents the time at which this via-point is traversed.

A set *X_m_* consisting of 251 minimum jerk trajectories were generated, corresponding to: (a) 25 different spatio-temporal constraints for each of 10 different values of *N*; and (b) one movement without any via-points. The sampling period of this problem was set to *δt_s_* = 0.001*s*, resulting in 1000 sample points per movement.

The smoothness of the movements in *X_m_* were analyzed using 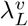 and 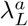. The scatter plot of 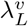 versus 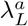 is shown in Fig. 10, which shows that the two measures are fairly well correlated, but are not isomorphic. This implies that the two measures have slightly different structures, and one cannot be exactly obtained from the other.

**Figure 10:**
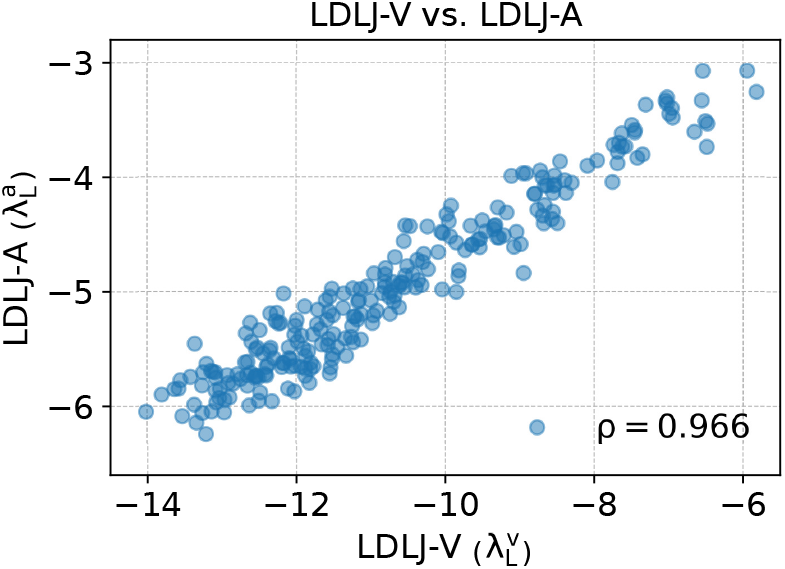
Scatter plot of 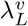 and 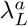 for a set of movements with varying number of submovements. The scatter plot shows that there is a fair correlation between the smoothness values reported by the two movements on the same movement data, but there is no one-to-one relationship.

## Appendix C SPARC on tranlsational velocity and acceleration data

The SPARC measure was also applied to the simulated data (from Section 7) to evaluate the measure’s consistency when estimated from movement velocity or acceleration (Fig. 11). Similar to Balasubramanian et al. (2015), there was a reasonably consistent response when using velocity data (Fig. 11(a)). On the other hand, when movement acceleration is used (Fig. 11(b)), it does not result in a consistent response (i.e., a monotonic response to change in inter-submovement interval).

**Figure 11:**
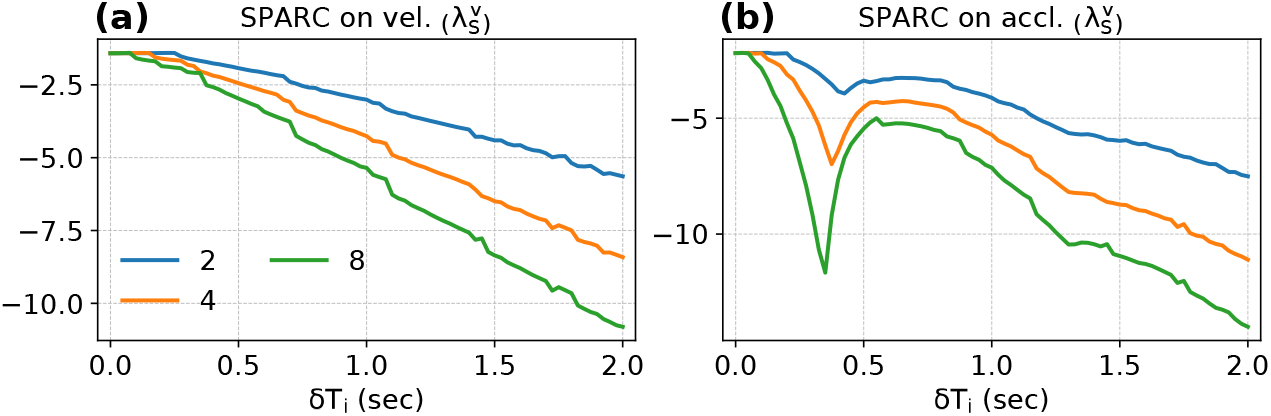
Consistency of 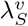 applied to (a) movement velocity and (b) movement acceleration.

This difference in behavior is due to the origins of SPARC: it was inspired by the idea of movement planning and execution by combining submovements. SPARC measures the level of dispersion of a signal in time (e.g., relative periods of movement and movement arrest). This measure of temporal dispersion acts as a reasonable measure of smoothness only when it is applied on the velocity, because velocity dispersion is strongly related to movement intermittency. Temporal dispersion in all other kinematic variables do not provide a unique measure of intermittency.

## Appendix D Calculating magnitude of linear jerk from IMU signals without rotation estimation

The derivative of the accelerometer signal 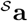 is given by (from Eqs. (3) and (4)):

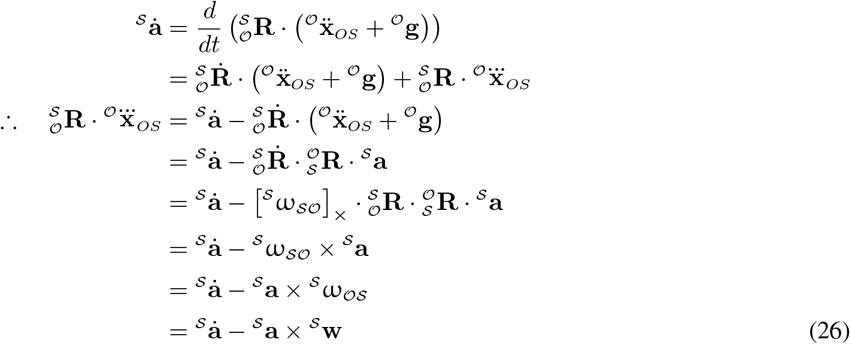

where the matrix 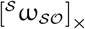 is skew-symmetric.

Because the operator ||*·*||_2_ is rotation invariant, the magnitude of the linear jerk is given by:

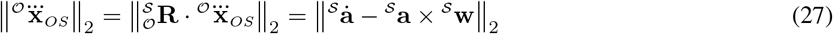

It is important to note that, in practice, the term 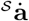 needs to be computed from numerical differentiation. Subtracting the correction term 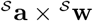, which is derived from the raw IMU signals, is affected by misalignment in time and different sampling of peaks. In simulated data, we found that, in some cases, this can lead to very erroneous estimation of the translational jerk. Further investigation is needed as to general recommendations that can be provided.

### Calculating magnitude of angular jerk from IMU signals without rotation estimation

Similar to translational jerk, one can compute the absolute angular jerk directly from IMU data. The second derivative of the gyroscope signal 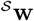 is given by:

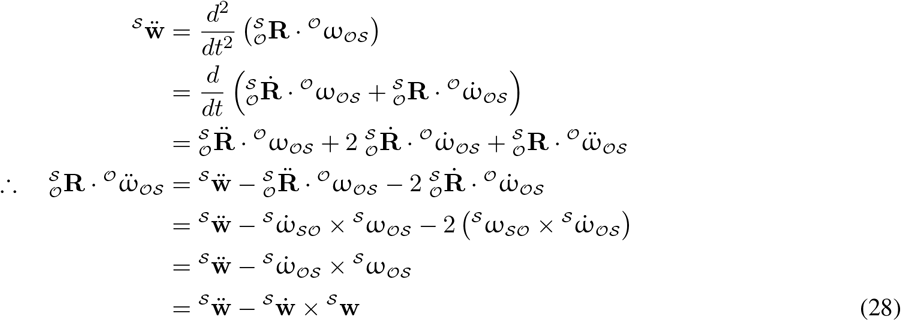

Thus, the magnitude of the absolute angular jerk is given by:

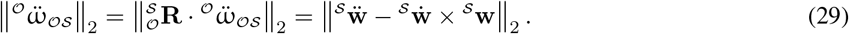

In practice, the terms 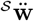 and 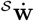 would need to be computed from numerical differentiation and may lead to erroneous estimations of the magnitude of angular jerk. Further investigation is needed as to general recommendations that can be provided.

